# Post-Assay Photogelation Enables Flow-Cytometric Sorting and Recovery of Bacterial Cocultures from Microfluidic Droplets

**DOI:** 10.64898/2026.08.27.747404

**Authors:** Ibraheem Alshareedah, Katelyn M. Green, Sang-Min Shin, Ramesh K. Jha, Anand Kumar

**Affiliations:** Microbial and Biome Sciences, Bioscience Division, Los Alamos National Laboratory, Los Alamos, New Mexico, USA; Nuclear and Radiochemistry, Chemistry Division, Los Alamos National Laboratory, Los Alamos, New Mexico, USA

## Abstract

High-throughput droplet microfluidics can compartmentalize bacterial interactions, but recovering droplets displaying phenotypes of interest often requires custom fluorescence-activated droplet-sorting instrumentation. Here, we introduce post-assay photogelation to decouple the material requirements of bacterial coculture from those of commercial flow sorting. Bacteria are cocultured in initially aqueous water-in-oil droplets containing photoreactive polymer precursors. After interaction phenotypes develop, ultraviolet exposure converts the droplets into mechanically stable hydrogel particles that can be transferred to an aqueous carrier and sorted using a commercial benchtop cell sorter. The sorted particles can subsequently be degraded enzymatically to release the encapsulated bacteria. We show that the timing of gelation alters bacterial growth and spatial distribution within droplets, with post-assay gelation supporting greater and more uniformly distributed growth than culture in preformed hydrogels. Using two fluorescent bead-encoded hydrogel-particle populations, we demonstrate sorting to greater than 99% purity. As an end-to-end demonstration, we cocultured sfGFP-expressing *Escherichia coli* Nissle 1917 with a cultured human nasal bacterial community and found that *E. coli* Nissle became the predominant detectable population under the tested conditions with possible inhibition of the cultured nasal bacteriome. This liquid-to-solid transition provides an accessible interface between aqueous bacterial droplet assays, commercial particle sorting, and downstream microbial analysis.

## Introduction

Complex bacterial communities, or ‘bacteriomes’, reside at multiple sites in and on the human body and profoundly influence immunity, health, and disease [1–11]. Perturbations of these communities have been associated with diseases affecting the lungs, gut, cervix, and other tissues[12–15]. Since key bacterial species can enhance physiological functions, preserve microbial homeostasis, and limit pathological deterioration[16], identifying important members of the bacteriome as well as key bacterial interactions may support the development of new microbiome-based therapies[17, 18]. Despite their importance, systematic study of interbacterial interactions remains difficult. Traditional methods that examine pairwise interactions or a small subset of model organisms provide mechanistic insights but scale poorly across the large number of possible strain combinations[19, 20]. Furthermore, interaction outcomes depend on strain identity, spatial organization[21], and environmental stress[22], making it difficult to extrapolate across contexts. Therefore, scalable platforms that can rapidly scan many bacterial combinations under controlled conditions are needed to facilitate the development of microbiome-based interventions to improve human health.

Droplet microfluidics has become widely adopted for high-throughput analysis of cells[23–25]. Its ability to miniaturize cellular assays has enabled many applications in single-cell sequencing[26], single-cell transcriptomics[27], discovery of antibiotic-resistant strains[28, 29], and bacterial cultivation[30]. Studies using droplet microfluidics to study bacterial interactions have focused on proof-of-concept cases using a few types of bacteria[31–34]. In fact, one study that used droplet microfluidics for cocultivation of bacteria highlighted the need for droplet sorting to enhance the throughput and robustness of their suggested pipeline[34]. However, droplet sorting often requires specialized techniques such as Fluorescence Activated Droplet Sorting (FADS)[35, 36]. FADS is commonly performed using custom-built instruments that rely on dielectric sorting, requiring expertise in optics and engineering[37].

Efforts have been made to develop alternative methods that modify droplet formulations to be compatible with conventional flow cytometry. This is an attractive approach because flow cytometers are widely available in research labs and institutional core facilities. For example, double emulsions have shown good compatibility with conventional flow cytometry, even though they require specialized microfluidic designs and coatings to allow consistent production of double emulsions[38]. Bacterial encapsulation in hydrogel particles also provides a strategy to tweak droplet microfluidics to be compatible with flow cytometry[38]. However, hydrogels can hinder bacterial growth and interactions due to their viscoelastic matrix[39]. Furthermore, common hydrogel formulations such as agarose can be digested by bacteria and become leaky, limiting the encapsulation time [40]. In short, *an accessible droplet microfluidic formulation that combines aqueous bacterial coculture, temporally controlled stabilization, conventional flow-cytometric sorting, and recovery of viable bacteria remains lacking*.

In this work, we present a highly tunable platform to co-encapsulate bacterial species in droplets, coculture them in aqueous media, convert them to a hydrogel post-culture using photopolymerization, sort them using a flow cytometer, and recover bacteria from sorted droplets via enzymatic degradation. To achieve post-assay photogelation, we use commercial synthetic polymers that are crosslinked by light to decouple bacterial culture from hydrogel formation. We evaluate PEG- and dextran-based formulations, identify a dextran-based degradable system that supports bacterial culture, and demonstrate efficient sorting using a commercial cell sorter. As an end-to-end validation, we coculture the fluorescent probiotic bacterium *E. coli* Nissle 1917 (EcN) with a cultured nasal bacteriome sample obtained commercially. Droplet sorting and subsequent recovery show that EcN apparently inhibited the growth of the nasal bacteriome in droplets under the tested conditions. Overall, this platform provides an accessible framework for coupling aqueous droplet bacterial coculture with commercial particle sorting and downstream microbial analysis via a temporally controlled liquid-to-gel transition.

## Results

### Post-assay photogelation preserves aqueous bacterial culture while enabling particle stabilization

Our platform is centered around five basic steps: encapsulation, incubation, gelation, sorting, and retrieval via degradation (**Fig. 1**). The first step is encapsulating bacterial species in microfluidic droplets containing a hydrogel precursor solution that can be crosslinked through UV light. We chose a norbornene-functionalized 8-arm polyethylene glycol (PEG) that has a molecular weight of 10,000 Daltons (**Fig. 2a**). For the crosslinker, we chose PEG dithiol (SH-PEG-SH) with a molecular weight of 1000 Daltons (**Fig. 2a**). Note that this crosslinker is not degradable; however, it serves as an initial choice to optimize the encapsulation and culture. The norbornene reacts with thiol groups on the crosslinker upon exposure to UV light only when there is a photoinitiator present. The photoinitiator used in this work is lithium phenyl-2,4,6-trimethylbenzoylphosphinate (LAP). We used a previously engineered EcN that expresses sfGFP as the bacterial species for initial platform development. We flowed EcN along with the gel precursor solution containing 7% (wt/vol) 8-arm PEG and 14 mM SH-PEG-SH in the presence of 0.05% (wt/vol) LAP into a microfluidic droplet generator with HFE7500 oil and a surfactant. This setup generated monodisperse water-in-oil (w/o) droplets with a diameter of 30 ± 1 µm and containing a small number of EcN cells (**Fig. 2b&c**).

**Figure 1:**
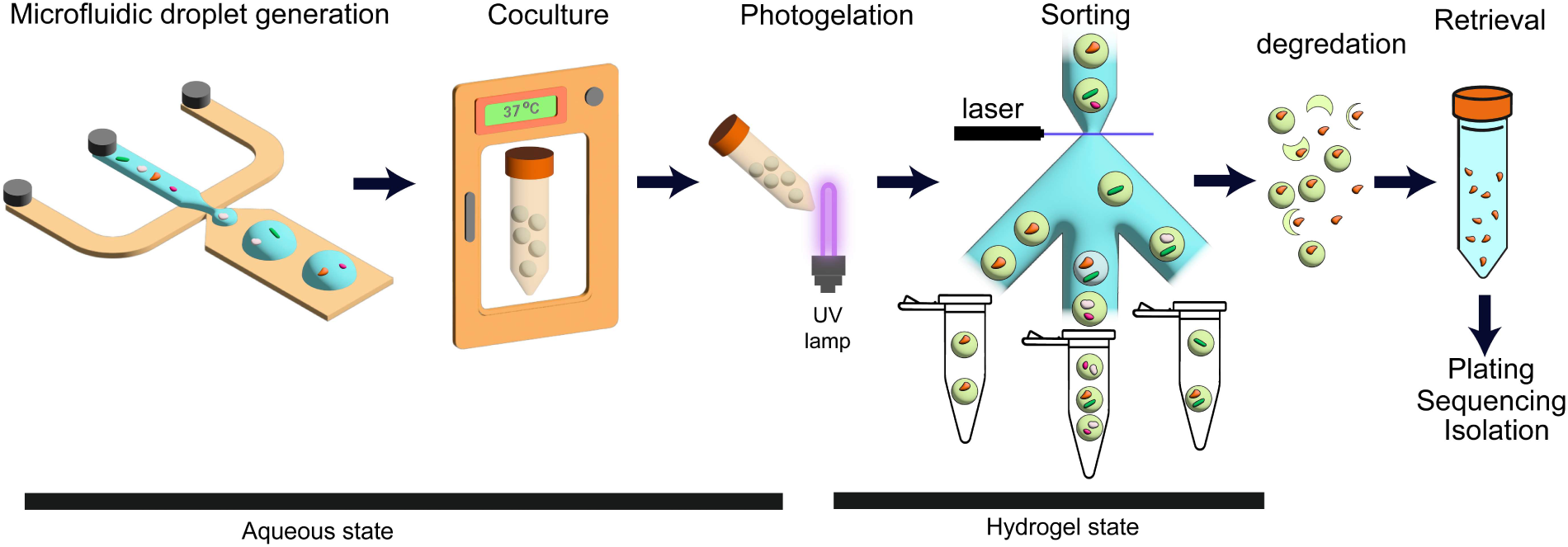
A droplet microfluidic platform for encapsulation, culture, sorting, and retrieval of bacterial cells.

**Figure 2:**
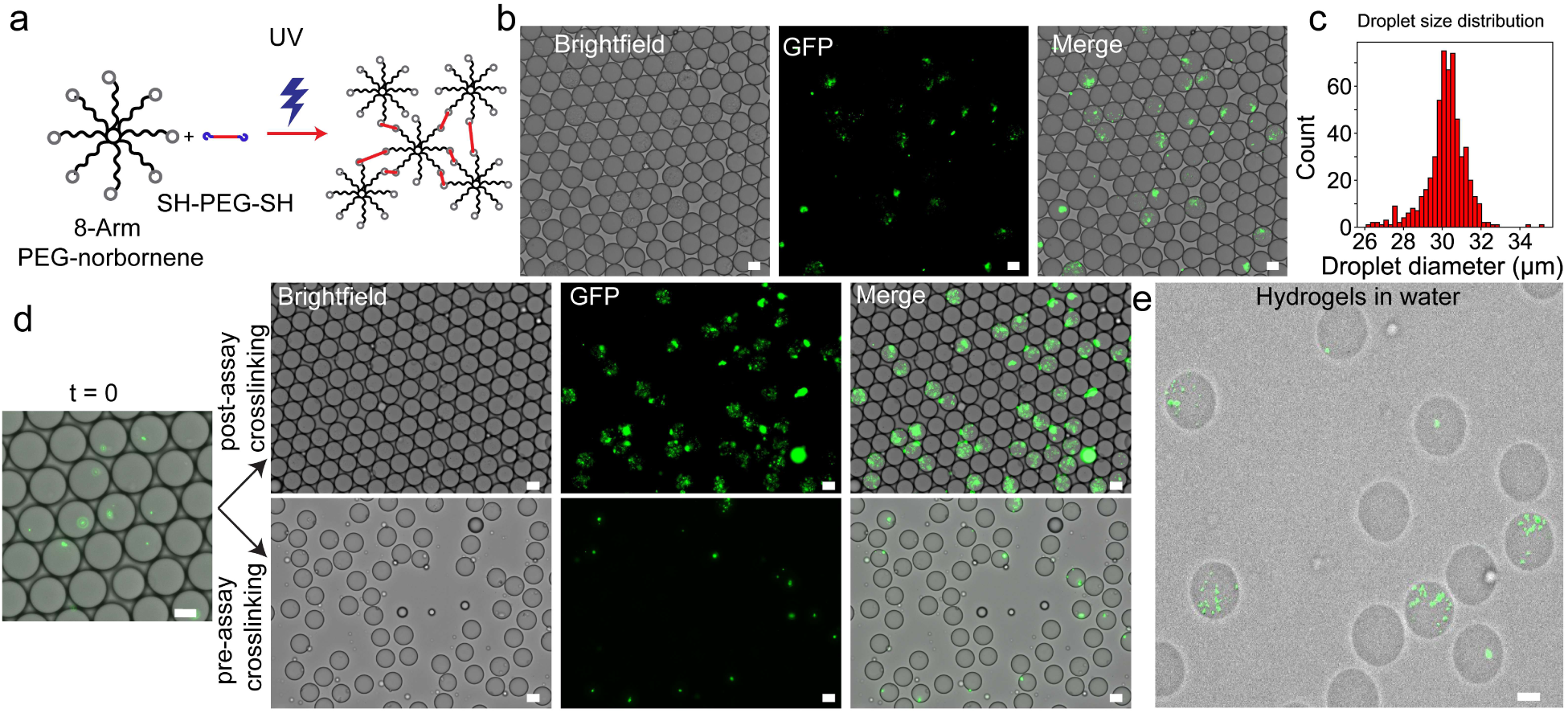
PEG hydrogel formulation with post-assay photogelation to culture bacterial cells. **(a)** Diagram showing the process of UV crosslinking of 8-arm PEG-norbornene and PEG dithiol as a crosslinker. **(b)** Brightfield and fluorescence images of GFP-expressing EcN cells encapsulated in microfluidic droplets. **(c)** Histogram of droplet diameters as measured by microscopy. **(d)** Brightfield and fluorescence images of droplets encapsulating EcN cells along with gel precursor solutions at t=0 and at t=18 hours in two conditions: pre-assay crosslinking (crosslinked right after encapsulation), and post-assay photogelation (crosslinked after 18 hours of growth). **(e)** EcN-laden hydrogels after transferring them to an aqueous carrier phase for sorting. All scale bars represent 20 µm.

We incubated the bacteria-laden droplets under two conditions: pre-assay and post-assay photogelation. In the first condition, we crosslinked the droplets via UV light (transforming them to a hydrogel) immediately after encapsulation such that the bacteria grow within the hydrogel droplets for 18 hours (pre-assay crosslinking, **Fig. 2d**). In the second condition, the bacteria grow in aqueous media for 18 hours before UV light-induced crosslinking (post-assay photogelation, **Fig. 2d**). We found that EcN grew significantly more in the aqueous media, and its growth was homogeneously distributed in the droplets (**Fig. 2d**). Conversely, EcN cells growing in the hydrogel particles formed small, dense, and bright fluorescent spheres (microcolonies, **Fig. 2d**). This shows that post-assay photogelation is preferred for measuring EcN dynamics in droplets, especially since droplet sorting relies on fluorescence and would distinguish different growth conditions more readily if the fluorescence signal is spread through the droplet as opposed to a dense signal from an EcN aggregate that is likely to saturate the fluorescence detector prematurely. It is also reported elsewhere that bacterial growth is much slower in a hydrogel compared to aqueous media[41]. Therefore, post-assay photogelation allows for aqueous culture that can be more comparable to classical bacterial culture, underscoring the importance of controlled UV-triggered gelation as opposed to chemical or temperature-based crosslinking that occurs at an uncontrolled time. Importantly, gelation allowed us to transfer the droplets from the oil carrier phase to an aqueous carrier phase to enable particle sorting, since oil is not compatible with flow cytometers due to its immiscibility with aqueous sheath fluids. After extraction, we obtained stable bacteria-carrying hydrogel particles that can be sorted using conventional flow cytometers (**Fig. 2e**).

### Crosslinker compatibility motivates the development of a degradable dextran formulation

High-throughput microdroplet studies of bacterial interactions require the ability to retrieve bacterial species of interest from droplets or hydrogels for further analysis, including sequencing and re-propagation. To do that, the hydrogel matrix must be degradable. The most convenient way to degrade hydrogels without exposing them to harsh conditions is enzymatic degradation. A common strategy is to use peptide crosslinkers that are cleavable by a sequence-specific protease. In studies involving human cell culture in hydrogels, MMP-cleavable peptides are used, as these can be cleaved by proteinases that remodel the extracellular matrix[42]. We therefore acquired a peptide that contains an MMP-cleavable motif and is flanked by two cysteine residues (thiols) that can crosslink our PEG-norbornene-based hydrogel (**Fig. 3a**). The peptide sequence is GCRDGPQGIAGQDRCG (henceforth referred to as the GPQ peptide). This GPQ peptide has been used previously to create degradable hydrogels for human cell culture that can be digested by commercial collagenase mixtures[42]. We tested the hydrogel degradability in bulk by preparing a gel precursor solution containing 10% PEG-NB and 20 mM GPQ peptide with 0.1% (wt/vol) LAP (photoinitiator). We then deposited 5 µL of the gel precursor solution into four wells and exposed them to UV light for gelation (**Fig. 3b**). The hydrogel also contained a fluorescent Dextran (Dextran-Texas Red, MW 70,000 Da) for visualization. The hydrogel was then submerged in PBS at different collagenase concentrations. Adding 5 mg/mL collagenase degraded the gel within 5 minutes, evidenced by the dispersion of the trapped fluorescent dextran molecules into the solution (**Fig. 3b**). At 1 mg/mL collagenase, the degradation took 15 minutes to completely remove the submerged hydrogel (**Fig. 3b**). Overall, our data confirms that crosslinking PEG with the GPQ peptide offers a viable gel that can be degraded by collagenase mixtures.

**Figure 3:**
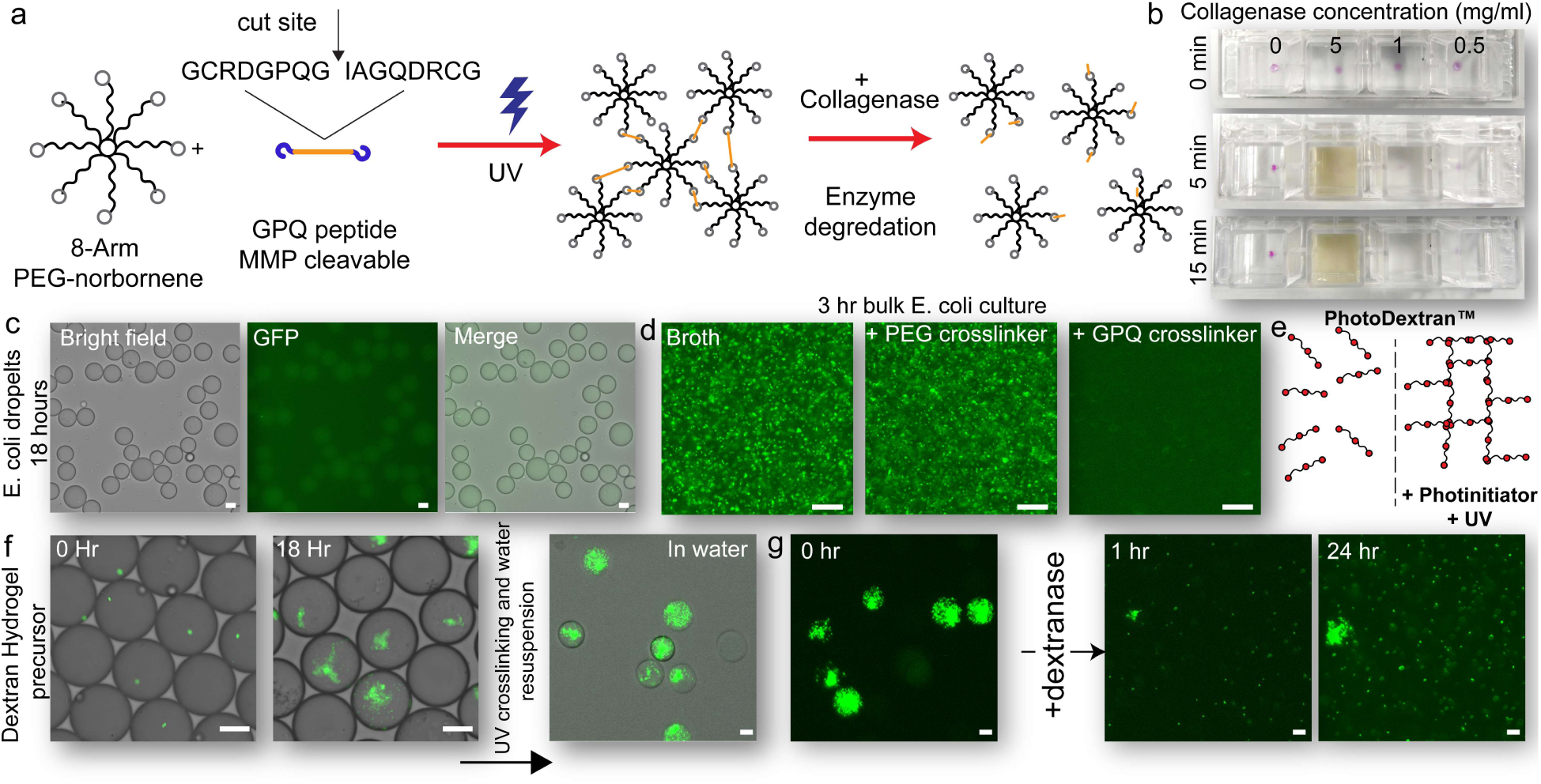
Enzymatic degradation of hydrogels and bacteria recovery. **(a)** A diagram showing the GPQ peptide as a crosslinker that can be cleaved by a collagenase enzyme mix. **(b)** Photographs showing hydrogels (pink drops) degraded by collagenase over time. **(c)** Brightfield and GFP images of droplets encapsulating EcN after 18 hours and containing GPQ as a crosslinker. **(d)** Fluorescence images showing the bulk culture of EcN in broth, broth with 14 mM PEG dithiol, and broth with 14 mM GPQ peptide. **(e)** Diagram showing photodextran, a dextran molecule carrying methacrylate groups along its linear chain, which enables UV-induced crosslinking. **(f)** Brightfield and GFP overlay images showing EcN growth in dextran hydrogel precursor droplets and crosslinked dextran hydrogel particles carrying grown EcN cultures. **(g)** Brightfield and GFP images showing the degradation of dextran hydrogels by dextranase and the release of EcN cells. All scale bars represent 20 µm.

To test whether our degradable hydrogel formulation can support bacterial growth, we added EcN cells to a gel precursor solution containing 7% 8-arm PEG-NB and 14 mM GPQ peptide in the presence of 0.05% (wt/vol) LAP. The solution was flowed into a microfluidic droplet generator. The EcN-encapsulating droplets were then placed in an incubator at 37 °C for growth. Counterintuitively, after 18 hours, fluorescence imaging revealed that all droplets in our sample were devoid of any fluorescent bacteria, suggesting no growth of EcN in droplets (**Fig. 3c**). We suspected that the GPQ peptide may have negative effects on the viability of EcN cells. We then prepared bulk EcN cultures in LB broth supplemented with either 14 mM of PEG dithiol or 14 mM of GPQ peptide, as well as plain LB broth culture as a control. Remarkably, the EcN culture containing the GPQ peptide showed no detectable bacteria, while the control (broth only) and the PEG dithiol culture showed extensive and similar growth (**Fig. 3d**). This led us to conclude that the GPQ peptide has inhibitory effects on EcN bacterial growth. We reason that the high concentration of the peptide (∼14 mM) may contribute to its EcN inhibition capacity. Overall, even though GPQ peptides are excellent crosslinkers for hydrogels supporting human cell culture[43], their use with bacteria may be detrimental in some cases, and alternatives should be sought.

To overcome the obstacle of GPQ toxicity without engineering a new degradable crosslinker sequence, we switched our hydrogel base polymer from PEG to dextran. Unlike PEG, dextran itself can be degraded by an enzyme called dextranase that is produced by some bacterial species. The advantage of dextran over other biopolymers is that it can be functionalized readily. We used a commercially available dextran-methacrylate (photodextran, M.W. 40,000) that can be crosslinked through the methacrylate groups upon exposure to UV light (**Fig. 3e**). Similar to the thiol-ene reaction, methacrylate crosslinking requires a photoinitiator such as LAP. We used 15% photodextran with 0.05% (wt/vol) LAP as our gel precursor solution. We added EcN to the mixture and generated droplets using our microfluidic droplet generator. After 18 hours of incubation of the droplet suspension, we found that EcN grew sufficiently in these droplets (**Fig. 3f**). We then crosslinked the droplets by exposing them to UV light and successfully transferred the hydrogels to water, confirming their stability and suitability for downstream analysis (**Fig. 3f**). We note that some bacterial species do produce dextranase[44, 45]; this possibility must be considered when maintaining bacterial cultures for extended periods. To confirm the recoverability of bacteria, we added dextranase to the hydrogel suspension and observed significant degradation of photodextran particles and the retrieval of EcN cells in solution (**Fig. 3g**).

### Commercial flow sorting of dextran hydrogel particles

To demonstrate sorting compatibility, we used a microfluidic droplet generator to generate two separate populations of dextran hydrogel particles (**Fig. 4a**). The first population encapsulated a large number of yellow-green carboxylate-modified fluorescent polystyrene beads with a nominal diameter of 200 nm. The second population of hydrogel droplets contained red fluorescent beads with the same diameter and surface modification (**Fig. 4a**). The carboxylate-modified surfaces were chosen so that the beads remain dispersed and do not accumulate at the water-oil interface during droplet generation. The two populations were then mixed to generate a pre-sort sample containing both green and red hydrogel droplets at an approximate ratio of 1:2 (green:red, **Fig. 4b**). We then flowed the pre-sort mixed hydrogel suspension into a WOLF G2 cell sorting instrument (NanoCellect) that relies on a piezoelectric pump to alter the fluid flow into sorting channels using multiple trigger options. To detect hydrogels, we relied on the backscattering of the hydrogel particles. **Figure 4c** shows the events detected by the flow cytometer using backscattering (y-axis) and green fluorescence signal (x-axis) excited by a 488 nm laser. As expected, we were able to see two populations of high-scattering particles (Log(BSC)>6) with two distinct green fluorescence populations (**Fig. 4c**). The low fluorescence events (FL∼10^3^) represent oil satellite droplets in the sample (**Fig. 4c&d**). We then selected two rectangular gates at Log(BSC) values of ∼6 and low and high green fluorescence signal (**Fig. 4c**). The lower green fluorescence population was expected to contain the red-fluorescent hydrogel particles (gate red, **Fig. 4c&d**). The higher green fluorescence events were expected to be the green-fluorescent hydrogel particles (gate green, **Fig. 4c&d**). Over the course of 1 hour, the instrument detected ∼50,000 events and sorted ∼12,000 hydrogels in the red gate and ∼7,000 hydrogel droplets in the green gate. Next, we took the sorted samples (green and red gates) and concentrated them via centrifugation. Subsequently, we captured multiple brightfield and fluorescence images of the sorted samples as well as the pre-sort sample. We observed that the sorted-green samples contained almost exclusively green-fluorescent hydrogels (**Fig. 4e**). On the other hand, the sorted-red sample contained red-fluorescent hydrogels (**Fig. 4f**). Image analysis was then used to count the red and green hydrogel populations in each of the three samples: pre-sort, sorted-green, and sorted-red. After analyzing thousands of hydrogel particles, we calculated the purity of each of the three samples. The pre-sort sample contained approximately 32% green hydrogels and 68% red hydrogels (**Fig. 4g**). After sorting, the sorted-green sample contained ∼99.9% green hydrogels while the sorted-red sample contained 0.3% green hydrogels (**Fig. 4g**). Based on this, we conclude that our recoverable dextran hydrogel particles can be sorted efficiently with conventional cell-sorter instruments without the need for fluorescence-activated droplet sorting technology (FADS).

**Figure 4:**
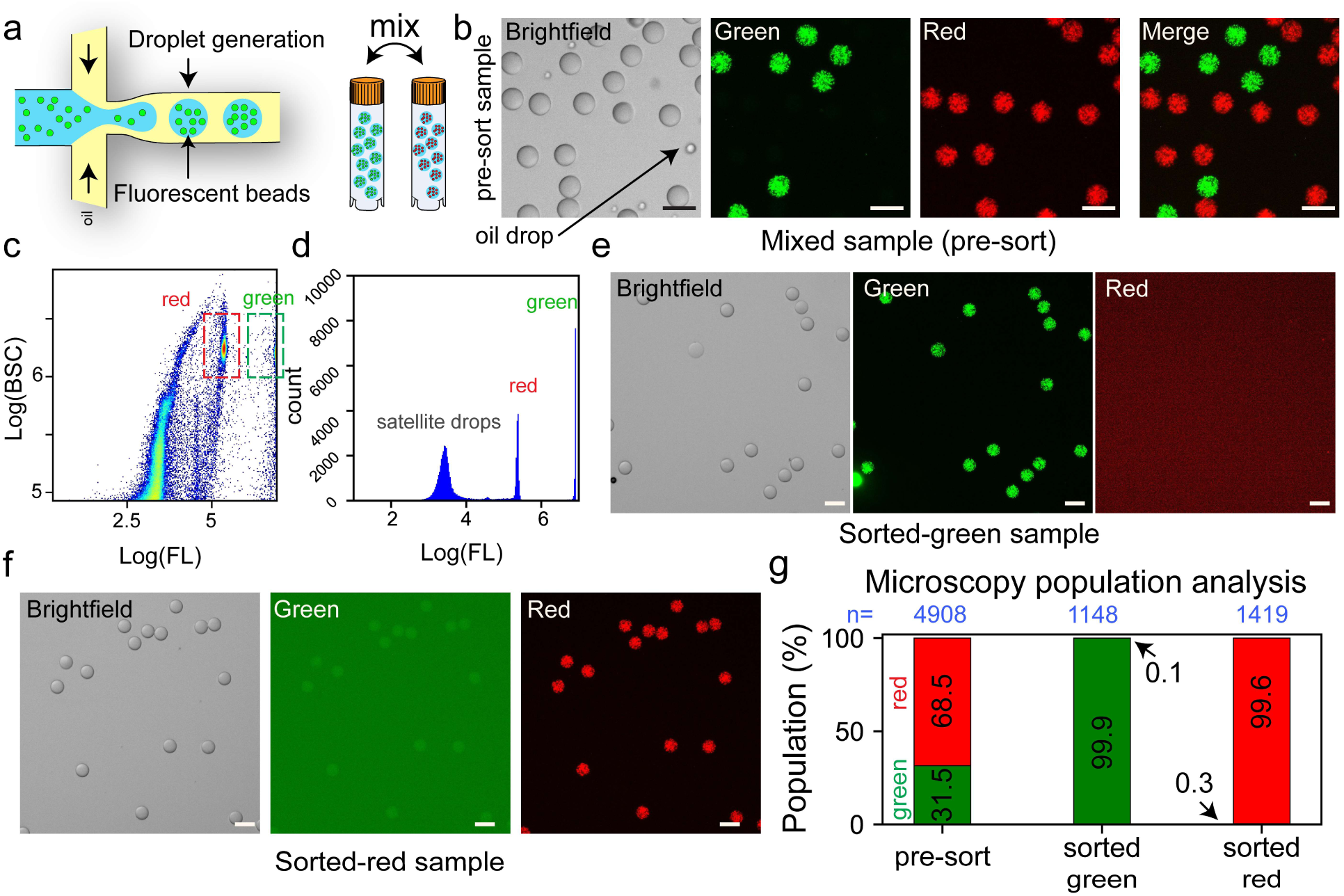
Droplet sorting of dextran hydrogel particles via a conventional cell sorter. **(a)** Schematic diagram of a microfluidic droplet generator to create two hydrogel samples encapsulating 200 nm fluorescent beads of two colors (green and red). These two samples are then mixed to create the pre-sort mixed population samples for hydrogel sorting tests. **(b)** Brightfield and fluorescence images of the pre-sort sample containing green and red hydrogels. **(c)** Flow cytometry heat map plotting backscattering value versus the fluorescence reading of the pre-sort sample excited with a 488 nm laser. The selected sorting gates are represented as green and red dashed rectangles. The unsorted events represent satellite oil droplets and contaminants. Two-way sorting was performed into the green and red channels while satellite droplets were passed through to the waste channel. **(d)** Histogram of the fluorescence readings of the hydrogels as measured by the cell sorter showing three distinct populations. **(e)** Brightfield and fluorescence images of the sorted hydrogels through the green gate. **(f)** Brightfield and fluorescence images of the sorted hydrogels through the red gate. **(g)** Microscopy-based population analysis of the sorted samples and the original pre-sort samples. This analysis was done on multiple images of these samples. Green and red droplet populations are quantified and plotted as a stacked bar graph. The contrast was enhanced for the red channel in sorted-green and the green channel in sorted-red for visualization purposes. The number of droplets analyzed is indicated above each bar. All scale bars represent 50 µm.

### End-to-end workflow demonstration using EcN and a cultured nasal bacterial community

In this section, we describe the application of our developed microfluidic droplet pipeline to study the interactions between EcN and a cultured nasal bacteriome sample. The purpose of this section is to show the feasibility and robustness of our droplet formulation in encapsulation, incubation, sorting, and retrieval of bacterial cocultures for screening bacteriome-target bacterial interactions. Briefly, we aerobically cultured and isolated the nasal bacteriome population from healthy human nasal swab samples obtained commercially (iSpecimen). The nasal bacteriome is relatively simple, exhibiting lower diversity and abundance than more complex microbial communities such as the gut microbiome[46]. This reduced complexity makes it a useful model for studying microbial interactions and developing experimental approaches before extending them to more complex microbiome ecosystems. We mixed EcN and bacteriome cultures at equivalent OD values in a solution containing 15% (wt/vol) photodextran and 0.05% (wt/vol) LAP and injected the solution into a microfluidic droplet generator chip. After droplet generation, we confirmed that most droplets encapsulated both EcN cells and nasal bacteriome cells (**Fig. 5a**). The droplet suspension was then incubated at 37 °C for 18 hours. Upon imaging, we found that some droplets contained only EcN culture with no sign of visible non-fluorescent bacteria (**Fig. 5b&c**), while other droplets had exclusively bacteriome cells with no detectable EcN (**Fig. 5b&c**). We did not find examples of droplets that contained both EcN and bacteriome together that were detectable by microscopy, even though we were able to see that the majority of the droplets at t=0 contained both EcN cells and nonfluorescent bacterial cells that belong to the bacteriome sample (**Fig. 5a**). We successfully crosslinked the droplets and transferred them to an aqueous carrier phase (**Fig. 5d**). We then sorted the hydrogels containing EcN and nasal bacteriome and separated fluorescent from nonfluorescent hydrogels (**Fig. 5e&f**). The fluorescent gels were then digested with dextranase to release the encapsulated bacteria. Upon analyzing the released bacteria with flow cytometry, we only found a single population with high fluorescence and low scattering that corresponds to EcN (**Fig. 5g**). This indicates that most of the hydrogels with detectable EcN cultures showed no detectable growth of nasal bacteriome cells. This is not a consequence of EcN outgrowing bacteriome samples since in droplets devoid of EcN, bacteriome populations grew substantially within the same time frame (**Fig. 5b**). Therefore, there may be an apparent inhibitory effect of EcN on the growth of this nasal bacteriome population.

**Figure 5:**
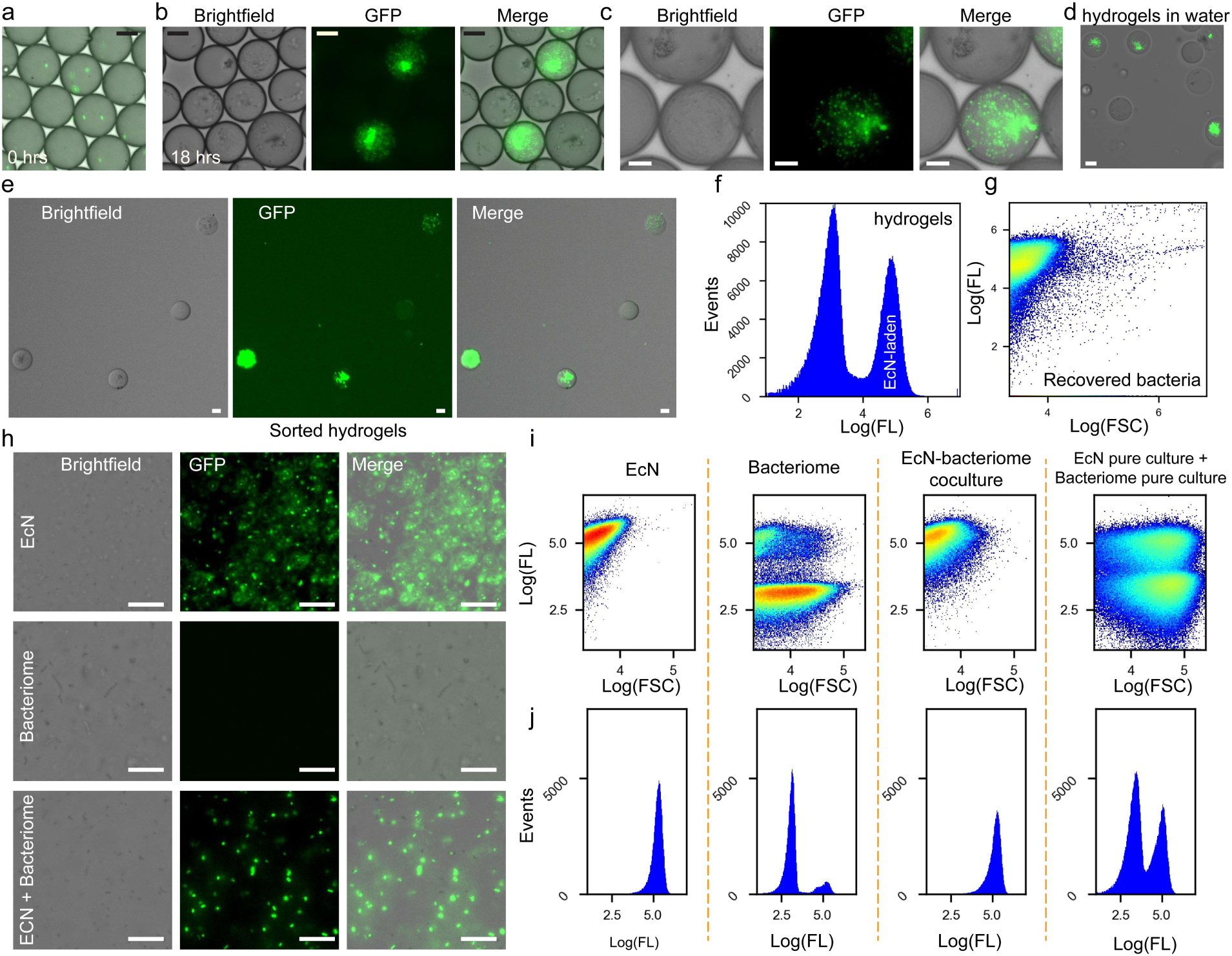
Exploring the interactions between EcN and nasal bacteriome via droplet microfluidics. **(a)** A brightfield and GFP image of droplets encapsulating EcN and nasal bacteriome cells immediately after droplet generation. **(b)** Brightfield and fluorescence images of droplets sustaining the culture of EcN and nasal bacteriome after 18 hours of incubation. **(c)** Zoomed-in brightfield and fluorescence images showing that droplets exclusively contain either the GFP-labeled EcN population or the nasal bacteriome population (nonfluorescent). **(d)** Brightfield and fluorescence images of EcN-bacteriome coculture hydrogels after crosslinking and transferring to water. **(e)** Brightfield and fluorescence images of the sorted EcN hydrogels. **(f)** Fluorescence histogram of hydrogels before sorting. **(g)** Forward scattering and fluorescence of bacteria released from sorted hydrogels showing one population of EcN and no detectable nonfluorescent population of the co-encapsulated nasal bacteriome. **(h)** Brightfield and fluorescence images of bulk cultures of EcN alone, bacteriome alone, and a coculture of EcN and bacteriome. These data show that the coculture of EcN and nasal bacteriome was dominated by EcN growth. **(i)** Flow cytometry plots of fluorescence versus forward scattering of bacterial cells analyzed using the flow cytometer for four samples: EcN culture, bacteriome culture, EcN-bacteriome coculture, and a mixture of separately grown pure EcN and bacteriome cultures. **(j)** Histogram of fluorescence distribution of the cells for the same samples shown in **(h) and (i)** as measured by the cell sorter. All scale bars represent 20 µm.

To further corroborate our observation, we performed bulk coculture experiments and found that while both EcN and bacteriome populations grew in TSB media when cultured separately, coculturing both types of cells produced a predominant EcN population after 18 hours of coculture with relatively few nonfluorescent bacteria visible under the microscope (**Fig. 5h**). We then prepared four samples of bulk cultures and flowed them into the cell sorter instrument with altered settings for detecting bacteria. Indeed, we saw that EcN cultures showed one population of cells that have high fluorescence and low scattering (**Fig. 5 i&j**). For the bacteriome sample, we saw a broader population having both low and high forward scattering as well as primarily low fluorescence **(Fig. 5 i&j)**. The broader scattering peak is consistent with multiple species and the presence of bacterial clusters as observed by microscopy. Interestingly, we saw a small population of cells emitting fluorescence signals that we attribute to autofluorescence of some nasal bacterial species.

We then ran the coculture sample that contained equivalent amounts of EcN and bacteriome at the start of the culture and saw that almost all the detected cells corresponded to lower scattering and higher fluorescence peaks consistent with EcN cells (**Fig. 5 i&j)**. As a control, we mixed grown bacteriome cultures and grown EcN cultures and immediately flowed them into the sorter and saw two clear populations corresponding to EcN (high fluorescence) and the bacteriome (low fluorescence, **Fig. 5 i&j**). This confirms that EcN apparently prevents the growth of this bacteriome population as qualitatively observed in microscopy and quantitatively observed in flow cytometry. This result is consistent with previous reports showing that EcN inhibits biofilm formation by *S. aureus* in vitro, and EcN culture supernatant (postbiotic) suppresses the growth of several bacterial species[47, 48]. Our findings therefore potentially generalize the effect of EcN to a broader community of species that constitute the cultured nasal bacteriome sample tested here. Downstream analysis and sequencing could identify the species present in this nasal bacteriome and determine whether any of them continue to grow in the presence of EcN. Overall, we used our platform to assess the compatibility of coculturing EcN cells with the nasal bacteriome and observed substantial inhibitory effects of EcN on the nasal bacteriome growth.

## Discussion and Conclusion

In this work, we present a microfluidic platform that decouples bacterial culture from hydrogel formation. Bacteria are first cultured in aqueous water-in-oil droplets; after the interaction phenotype has developed, the droplets are stabilized through photo-induced crosslinking, transferred to a water-based carrier, sorted with a conventional cell sorter, and enzymatically degraded to recover the encapsulated bacteria. The approach addresses a significant challenge in droplet-based bacterial screening: maintaining an aqueous microbe-friendly environment while producing particles with sufficient mechanical stability and compatibility with downstream flow cytometric sorting.

Current droplet sorting approaches have trade-offs between biological compatibility, accessibility, and recoverability. Fluorescence-activated droplet sorting requires custom optical, electronic, and microfluidic instrumentation[36, 49]. Double emulsions have been presented as a solution to increase compatibility with flow cytometry[38, 50]; however, they require specialized microfluidic chips with non-uniform hydrophobic-hydrophilic coatings and precise fluidic control. Hydrogel particles have been used for aqueous sorting[51–53], although pre-assay gelation may restrain bacterial growth due to hindered diffusion and stiff matrix mechanics. By introducing post-assay photogelation, our platform separates the biological assay from the material requirements for sorting. In other words, the aqueous droplets support bacterial growth and interactions, whereas the subsequent photo-induced crosslinking arrests the resulting phenotype in a mechanically stable and sortable particle.

Our comparison of the PEG and dextran hydrogel chemistries highlights the importance of designing materials specifically for microbial studies. While the PEG formulation with the MMP-degradable peptides has been used for human cell culture[54, 55], we observe that the tested GPQ peptide inhibited EcN growth. Therefore, the compatibility of MMP-sensitive crosslinking peptides with bacteria cannot be assumed. Switching to methacrylated dextran provided a UV-crosslinkable base polymer that supports bacterial culture, sorting, and recovery via the dextranase enzyme. This finding emphasizes that degradability alone is insufficient. The entire matrix base, crosslinking mechanism, and degradation chemistry must be compatible with the studied organism.

Using bead-loaded dextran hydrogels with different colors, we demonstrated efficient hydrogel sorting with the WOLF G2 cell sorter (NanoCellect) that utilizes gentle hydrodynamic pressure for sorting. We observed high purity of the sorted populations, indicating that our platform is compatible with conventional cell-sorting instruments without the need for FADS systems. However, it must be noted that this compatibility does not immediately translate into other flow cytometers. Air-jet-based sorting systems may expose hydrogels to different hydrodynamic pressures, sorting-junction geometries, and shear stresses that can hinder the sorting performance. Nevertheless, the proposed platform offers sufficient flexibility to be fine-tuned with other flow cytometers by adjusting the polymer weight, crosslinking density, UV exposure time, and photoinitiator concentration to encode the necessary material properties. Accordingly, compatibility with other flow cytometers should be optimized before deploying this method.

As a proof of concept, we use our platform to test the coculture compatibility between EcN and a cultured nasal bacteriome sample. We observed that EcN inhibited the growth of the nasal bacteriome in both droplets and bulk coculture. This observation is consistent with recent reports showing that EcN inhibits biofilm formation and for certain bacterial species such as *Staphylococcus aureus*[47, 48]. Notably, *Staphylococcus* is one of the predominant bacterial genera in the human nasal microbiome[56, 57]. Our findings suggest that the inhibitory activity of EcN may extend beyond *S. aureus*, suppressing the growth of multiple bacterial species and genera within the cultured nasal bacterial community under the tested conditions. However, a limitation of our study is that taxonomic identification before and after sorting was not performed. This is necessary to reveal which organisms are affected by the presence of EcN. Future work will focus on investigating these interactions from a microbiological perspective. Other limitations of this work include optimizing particle degradation, investigating the effect of UV exposure on the bacterial community tested, probing the effect of dextranase-producing bacteria on particle stability, and method validation for a large combinatorial screen rather than single-community-level proof of concept. Addressing these limitations will be important to deploy this method for full-scale microbial library screens.

In conclusion, we propose a controlled and recoverable droplet microfluidic workflow that combines aqueous bacterial coculture, post-assay photogelation, conventional cell sorting, and bacterial recovery via enzymatic degradation of the sorted particles. This approach eliminates the need for a custom-built droplet sorter, providing high-purity sorting of dextran-based hydrogels and enabling a preliminary analysis of the culture compatibility of EcN with a cultured nasal bacteriome sample. Our approach could provide an accessible framework for identifying key bacterial strains and interactions relevant to microbiome research and biotechnology.

## Materials and Methods

### Ethics statement

De-identified human nasal swab samples were obtained commercially from iSpecimen. Per vendor documentation, specimens were collected under applicable IRB/IEC or other appropriate ethics oversight, with informed consent, waiver of consent, or a non-human-subjects research determination as appropriate for the specimen source. The authors received only de-identified samples and associated non-identifying donor information.

### GFP-expressing EcN

A GFP-expressing EcN strain was constructed using sfpGLOpnpv6, a plasmid in which superfolder GFP (sfGFP) is expressed from a promoter requiring an exogenous small-molecule inducer[58]. To achieve constitutive GFP expression in EcN, the promoter of the native *katG* (encoding catalase-peroxidase) gene, together with a ribosome binding site, was synthesized (Twist Bioscience). This fragment was used to replace the existing promoter region of the *sfgfp* reporter gene via the ClaI and NheI sites, which also removed the gene encoding the cognate transcription factor, yielding pKG01. The *katG* promoter showed constitutive activity in EcN in the growth media background, enabling stable GFP expression during bacterial growth. pKG01 was introduced by electroporation (BTX ECM 630; 1.8 kV, 25 μF, 200 Ω, 0.1 cm cuvettes) into in-house-prepared EcN competent cells generated from mid-log-phase cultures using a chilled 300 mM sucrose protocol. Following recovery, transformants were selected on LB agar plates supplemented with carbenicillin (50 μg/mL). Plasmid DNA was isolated from positive transformants using a QIAprep Spin Miniprep Kit (Qiagen) and verified by Sanger sequencing to confirm the correct promoter substitution. Sequence-verified transformants were preserved in LB glycerol stocks (20% v/v) and stored at −80 °C. For routine maintenance, GFP-expressing EcN cultures were grown in LB medium supplemented with carbenicillin (50 μg/mL) to maintain plasmid selection. For all coculture and microdroplet encapsulation experiments, however, bacteria were cultured in antibiotic-free medium to avoid potential effects on cocultured bacterial species.

#### Nasal Bacteriome Culturing

Human nasal swab samples were procured from iSpecimen (commercial vendor), and per vendor documentation, donors had no respiratory infections. Samples were thawed and vortexed briefly to generate a uniform microbial suspension in transport medium. A 200 µL aliquot was plated onto three commonly used media: (1) tryptic soy agar (TSA), (2) brain heart infusion agar (BHI), and (3) TSA supplemented with 10% defibrinated sheep blood to retrieve diverse bacterial isolates supported by different media compositions. The inoculum was spread evenly using sterile disposable L-shaped spreaders, and plates were incubated aerobically at 37 °C for 48 hours. Well-separated colonies were picked and restreaked onto agar plates prior to subculturing in their respective broth media: tryptic soy broth (TSB), brain heart infusion broth (BHI), or TSB supplemented with 10% defibrinated sheep blood. Purified isolates were prepared as glycerol stocks and stored at −80°C. Since the maximum number of diverse colonies was obtained from TSA plates, all isolates were combined in TSB to generate a composite bacteriome culture and incubated aerobically for 24 hours. For the present study, the pooled nasal bacteriome was used to investigate interactions with EcN.

### Culturing EcN in aqueous PEG droplets

8-arm PEG Norbornene (M.W. 10,000) and PEG dithiol (SH-PEG-SH, M.W. 1000) were purchased from Creative PEGWorks and used without further purification. To prepare microfluidic droplets using gel precursor solution, we first made a solution containing 7 % (wt/vol) 8-arm PEG-NB, 14 mM PEG-dithiol, and 0.05 wt% LAP (Lithium phenyl-2,4,6-trimethylbenzoylphosphinate, Sigma-Aldrich). EcN was added to the solution at a final estimated OD_600_ of 0.02. The final solution was flowed into a microfluidic chip with cross-junction geometry that was custom designed to have a nozzle of 20 µm (uFluidics) as described in our previous work[59]. HFE7500 oil with 2 wt% 008-fluorosurfactant (RAN Biotechnologies) was used as the carrier phase. Flow rates were adjusted to produce droplets with diameters of ∼30-40 µm. The droplets were covered with anti-evaporation oil (Ibidi) and placed in a dark incubator at 37 °C for 18-24 hours.

After incubation, the droplet suspension was taken out and imaged under an EVOS M5000 microscope equipped with GFP, RFP, and Texas Red light cubes (Invitrogen). The droplet suspension was then exposed to 395 nm UV light (AmScope) for less than 60 seconds to activate thiol-ene crosslinking. After crosslinking, the hydrogel droplet suspension was separated from the anti-evaporation oil by careful pipetting and mixed with an equal volume of 20 vol% 1H,1H,2H,2H-Perfluoro-1-octanol (diluted in HFE7500 oil) to break the emulsion. After mixing with a pipette, the sample was centrifuged for 5 minutes at 2000 x g. This produced a pellet containing hydrogel droplets. The HFE7500 oil was removed by pipetting and discarded. A PBS solution with 0.05% Poloxamer 188 was then added to the pellet and vortexed to resuspend the hydrogels. The hydrogel droplets were washed twice with the 0.05% Poloxamer 188 solution using a 10 µm cell strainer (Pluriselect Mini) to remove residual oil and satellite droplets. The hydrogels were then stored at 4 °C for further processing and imaging.

### *E. coli* culture in degradable PEG hydrogels

To prepare degradable PEG hydrogels, we replaced PEG dithiol with the GPQ peptide (GCRDGPQGIAGQDRCG) that features a cleavable motif and two cysteine residues that can react with norbornene through their thiol groups. GPQ peptide was obtained from GenScript and used without further purification. The gel precursor solution was prepared at 7 % (wt/vol) 8-arm PEG Norbornene, 14 mM GPQ peptide, and 0.05% LAP.

To confirm the degradability of the gel, we first placed 5 µL of the gel precursor solution doped with dextran-Texas Red (M.W., 70,000) in four wells of an 8-well imaging chamber. The chamber was exposed to UV light for 60 seconds to harden the gels. Gels were then submerged in PBS containing variable concentrations of the collagenase enzyme mixture (Collagenase from *Clostridium histolyticum*, Sigma-Aldrich). We then took photographs of the 8-well chamber after 5 minutes and 15 minutes to assess the degradation of the gel (the disappearance of the red gel droplet at the bottom of the well).

To evaluate the compatibility of the degradable PEG hydrogel with *E. coli* growth, we added EcN to the gel precursor solution (7 % (wt/vol) 8-arm PEG Norbornene, 14 mM GPQ peptide, and 0.05% (wt/vol) LAP) to a final estimated OD_600_ of 0.02 and then flowed the solution into our microfluidic droplet generator. After droplet generation, the droplet suspension was covered with Ibidi anti-evaporation oil and placed in a dark incubator at 37 °C for 18 hours. Samples were then imaged to check for bacterial growth or lack thereof.

For bulk EcN culture experiments in the presence of crosslinkers, we prepared sterile LB medium. 5 µL of overnight-grown culture (containing ∼10^6^ cfu) of bacteria (EcN) was added to 2 mL of broth and placed in an incubator at 37 °C. Two additional samples were prepared by supplementing the broth separately with 14 mM GPQ peptide or 14 mM PEG-dithiol and simultaneously incubated at 37 °C. Imaging was done after three hours of incubation using an EVOS M5000 microscope.

### Bacterial culture in degradable dextran hydrogels

To prepare a degradable dextran hydrogel, we used methacrylated dextran (photodextran, Advanced BioMatrix) reconstituted in TSB media to support bacterial growth. Photodextran was mixed with LAP at a final concentration of 15 % (wt/vol) photodextran and 0.05 % (wt/vol) LAP. EcN was added to a final estimated OD of 0.02. The gel precursor solution was then flowed into our droplet generator as described before to produce droplets. The droplet suspension was then incubated at 37 °C for 18 hours. After that, droplets were exposed to UV light for less than 60 seconds to crosslink photodextran. The gelled droplets were then transferred to PBS as described earlier and stored at 4 °C.

For degradation experiments, dextranase (Dextranase from *Penicillium sp.*, Sigma Aldrich) was added to a final concentration of 15 U/mL to the hydrogel droplet suspension in PBS (pH 7.4) and incubated at 37 °C for 24 hours. We noted that complete degradation was not observed at 24 hours; we were able to see some gels still intact. However, the majority of the hydrogel droplets were degraded, and bacteria were released into the carrier phase. The released bacteria were then filtered using a 10 µm mini-cell strainer before flow cytometric analysis.

### Bead-laden hydrogel droplet sorting

For the bead-laden hydrogel sorting, we diluted yellow-green fluorescence beads that have carboxylate-functionalized surfaces (Invitrogen) in 10% (wt/vol) photodextran solution (Advanced BioMatrix) and 0.05% (wt/vol) LAP to a final bead concentration of 0.01% solids. The solution was introduced into a flow-focusing microfluidic chamber (uFluidics) to generate droplets with a mean diameter of 35 µm. The droplets were then exposed to UV light using a UV lamp to induce photo-crosslinking. Another population of droplets was prepared identically but with red-fluorescent beads (Invitrogen). The two samples were then mixed and filtered with a 10 µm cell strainer to remove oil satellite droplets.

Dextran hydrogels suspended in 0.05% Poloxamer 188 and 0.1 % BSA were analyzed and sorted using the WOLF G2 cell sorter (NanoCellect). WOLF G2 sorts cells using a four-way junction with a piezoelectric pump that applies pressure or suction to redirect the flow into its sorting channels. The WOLF cartridges are loaded into the instrument and primed using PBS as both sample and sheath fluids. A solution containing 1% BSA solution with 0.1% Poloxamer 188 was flowed for 15 minutes into the sample channel to ensure passivation and prevent the gels from sticking to the flow chamber surface. After alignment of the detection laser, green calibration beads (NanoCellect) were used to calibrate the instrument for bulk sorting. Hydrogels were first filtered with a 40 µm cell strainer to prevent clogging and remove any hydrogel aggregates. Hydrogels were then loaded into the sample reservoir after sufficient flushing and analyzed using various signals such as forward scattering, backscattering, and fluorescence. Sorting gates were selected based on the fluorescence signal to sort hydrogels. A typical hydrogel-sorting session will sort approximately 20,000 hydrogels in 60 minutes, depending on the concentration of the hydrogels in the input sample. The sorted hydrogels were then concentrated using a 10 µm mini-cell strainer or centrifugation. Subsequent analysis involved imaging the sorted samples via a fluorescence microscope (EVOS M5000). A custom Python code was used to count the number of green and red droplets in microscopy images by using thresholding to identify the droplets in both the green and red channels and subsequently calculate the distribution.

### Coculture of EcN and Nasal bacteriome in dextran hydrogels

A solution containing 15% photodextran (Advanced BioMatrix) with 0.05% LAP in TSB media was prepared. EcN and nasal bacteriome were each added to a final estimated OD of 0.02. The gel precursor solution was then flowed into a microfluidic droplet generator chip (uFluidics) to generate droplets ∼35-45 µm in diameter. Droplets were incubated at 37 °C for 18 hours. After incubation, droplets were crosslinked by UV light exposure for 60 seconds and transferred to an aqueous solution containing 0.01% Poloxamer 188 as described earlier. The droplet suspension was then analyzed via the WOLF G2 cell sorter, and fluorescent hydrogels (containing EcN) were sorted. The sorted sample was then degraded using dextranase (Dextranase from *Penicillium sp.*, Sigma-Aldrich). Next, the released bacteria were filtered using a 10 µm mini-cell strainer to remove the undegraded gels. The retrieved bacterial sample was then flowed into the cell sorter for flow cytometric analysis.

### Bulk coculture experiments

For EcN-nasal bacteriome bulk coculture experiments, three samples were prepared containing EcN, nasal bacteriome, and a mixture of EcN and nasal bacteriome suspended in sterile TSB media. The samples were incubated at 37 °C for 18 hours. After incubation, the samples were imaged using a fluorescence microscope (EVOS M5000).

To perform flow cytometric analysis, the three samples were filtered using a 10 µm mini-cell strainer and flowed into the cell sorter chip (WOLF G2, NanoCellect) consecutively with a 3-minute PBS flush between each sample run. A fourth sample was generated by mixing the grown pure EcN culture with the grown pure nasal bacteriome culture. The resulting mixture was immediately analyzed using the cell sorter to determine the scattering and fluorescence distributions.

### Plots and figures

All graphs and plots were generated using custom-made Python scripts. Artwork was generated using Adobe Illustrator.

## Acknowledgments

This work was supported by the U.S. Department of Energy (DOE) through Los Alamos National Laboratory, which is operated by Triad National Security, LLC, for the National Nuclear Security Administration of U.S. DOE (Contract No. 89233218CNA000001). The authors acknowledge support from the Los Alamos National Laboratory (LANL) Laboratory Directed Research and Development (LDRD) program. This work benefited from resources and reagents developed through the LDRD project (0240803ER: A Living Diagnostic–Therapeutic System), awarded to A.K. and R.K.J. We gratefully acknowledge Kartika Wardhani for her initial efforts in processing human nasal swab samples and isolating and banking nasal bacterial isolates under the supervision of A.K. The commercially-obtained nasal swab samples used in this study were processed as part of the ongoing LDRD project (20240092DR: The Next-Gen Approach to Rapidly Dissect Complex Host–Microbe Interactions), awarded to A.K. and colleagues to investigate the role of the nasal bacteriome in respiratory syncytial virus (RSV) infection. I.A. was supported by the Frederick Reines Fellowship (LDRD 20251143PRD1) at Los Alamos National Laboratory. LA-UR-26-25782.

## Data Availability

All data relevant to the conclusions of this manuscript are presented in the main text. All data are available from the authors upon reasonable request.

## References

1. Barlow GM, Yu A, Mathur R. Role of the gut microbiome in obesity and diabetes mellitus. Nutrition in clinical practice. 2015;30(6):787–97.

2. Blaser MJ. The microbiome revolution. The Journal of clinical investigation. 2014;124(10):4162–5.

3. Caporaso JG, Lauber CL, Costello EK, Berg-Lyons D, Gonzalez A, Stombaugh J, et al. Moving pictures of the human microbiome. Genome biology. 2011;12(5):R50.

4. Cullin N, Azevedo Antunes C, Straussman R, Stein-Thoeringer CK, Elinav E. Microbiome and cancer. Cancer Cell. 2021;39(10):1317–41. doi: 10.1016/j.ccell.2021.08.006.

5. de Vos WM, de Vos EA. Role of the intestinal microbiome in health and disease: from correlation to causation. Nutrition reviews. 2012;70(suppl_1):S45–S56.

6. De Vos WM, Tilg H, Van Hul M, Cani PD. Gut microbiome and health: mechanistic insights. Gut. 2022;71(5):1020–32.

7. Dohlman AB, Mendoza DA, Ding S, Gao M, Dressman H, Iliev ID, et al. The cancer microbiome atlas: a pan-cancer comparative analysis to distinguish tissue-resident microbiota from contaminants. Cell host & microbe. 2021;29(2):281–98. e5.

8. Durack J, Lynch SV. The gut microbiome: Relationships with disease and opportunities for therapy. Journal of experimental medicine. 2019;216(1):20–40.

9. Belizário J, Garay-Malpartida M, Faintuch J. Lung microbiome and origins of the respiratory diseases. Current Research in Immunology. 2023;4:100065.

10. Deo PN, Deshmukh R. Oral microbiome: Unveiling the fundamentals. Journal of oral and maxillofacial pathology. 2019;23(1):122–8.

11. Fan X, Alekseyenko AV, Wu J, Peters BA, Jacobs EJ, Gapstur SM, et al. Human oral microbiome and prospective risk for pancreatic cancer: a population-based nested case-control study. Gut. 2018;67(1):120–7.

12. Liu N-N, Ma Q, Ge Y, Yi C-X, Wei L-Q, Tan J-C, et al. Microbiome dysbiosis in lung cancer: from composition to therapy. NPJ precision oncology. 2020;4(1):33.

13. Liu W, Zhang R, Shu R, Yu J, Li H, Long H, et al. Study of the relationship between microbiome and colorectal cancer susceptibility using 16SrRNA sequencing. BioMed research international. 2020;2020(1):7828392.

14. Menees S, Chey W. The gut microbiome and irritable bowel syndrome. F1000Research. 2018;7:F1000 Faculty Rev-29.

15. Huang S-T, Chen J, Lian L-Y, Cai H-H, Zeng H-S, Zheng M, et al. Intratumoral levels and prognostic significance of Fusobacterium nucleatum in cervical carcinoma. Aging (Albany NY). 2020;12(22):23337.

16. Caballero-Flores G, Pickard JM, Núñez G. Microbiota-mediated colonization resistance: mechanisms and regulation. Nature Reviews Microbiology. 2023;21(6):347–60. doi: 10.1038/s41579-022-00833-7.

17. Henderson EA, Lukomski S, Boone BA. Emerging applications of cancer bacteriotherapy towards treatment of pancreatic cancer. Frontiers in Oncology. 2023;13:1217095.

18. Fan JY, Huang Y, Li Y, Muluh TA, Fu SZ, Wu JB. Bacteria in cancer therapy: A new generation of weapons. Cancer Medicine. 2022;11(23):4457–68.

19. Weiss AS, Burrichter AG, Durai Raj AC, von Strempel A, Meng C, Kleigrewe K, et al. In vitro interaction network of a synthetic gut bacterial community. The ISME journal. 2022;16(4):1095–109.

20. Gupta G, Ndiaye A, Filteau M. Leveraging experimental strategies to capture different dimensions of microbial interactions. Frontiers in Microbiology. 2021;12:700752.

21. Ackermann M, van Vliet S. Spatial self-organization of metabolism in microbial systems: a matter of enzymes and chemicals. Cell systems. 2023;14(2):98–108.

22. Hernandez DJ, David AS, Menges ES, Searcy CA, Afkhami ME. Environmental stress destabilizes microbial networks. The ISME journal. 2021;15(6):1722–34.

23. Dortaj H, Amani AM, Tayebi L, Azarpira N, Ghasemi Toudeshkchouei M, Hassanpour-Dehnavi A, et al. Droplet-based microfluidics: an efficient high-throughput portable system for cell encapsulation. Journal of Microencapsulation. 2024;41(6):479–501.

24. Yan W, Li X, Zhao D, Xie M, Li T, Qian L, et al. Advanced strategies in high-throughput droplet screening for enzyme engineering. Biosensors and Bioelectronics. 2024;248:115972.

25. Nan L, Zhang H, Weitz DA, Shum HC. Development and future of droplet microfluidics. Lab on a Chip. 2024;24(5):1135–53.

26. Hu B, Xu P, Ma L, Chen D, Wang J, Dai X, et al. One cell at a time: droplet-based microbial cultivation, screening and sequencing. Marine life science & technology. 2021;3(2):169–88.

27. Xu Z, Wang Y, Sheng K, Rosenthal R, Liu N, Hua X, et al. Droplet-based high-throughput single microbe RNA sequencing by smRandom-seq. Nature communications. 2023;14(1):5130.

28. Postek W, Garstecki P. Droplet microfluidics for high-throughput analysis of antibiotic susceptibility in bacterial cells and populations. Accounts of chemical research. 2022;55(5):605–15.

29. Hsieh K, Mach KE, Zhang P, Liao JC, Wang T-H. Combating antimicrobial resistance via single-cell diagnostic technologies powered by droplet microfluidics. Accounts of chemical research. 2021;55(2):123–33.

30. Ma H, Zhang Y, Shen R, Jia Y. Droplet-Based Microfluidics in Single-Bacterium Analysis: Advancements in Cultivation, Detection, and Application. Biosensors. 2025;15(8):535.

31. Park J, Kerner A, Burns MA, Lin XN. Microdroplet-enabled highly parallel co-cultivation of microbial communities. PloS one. 2011;6(2):e17019.

32. Jiang M-Z, Zhu H-Z, Zhou N, Liu C, Jiang C-Y, Wang Y, et al. Droplet microfluidics-based high-throughput bacterial cultivation for validation of taxon pairs in microbial co-occurrence networks. Scientific Reports. 2022;12(1):18145.

33. Jung S-Y, Kim KH, Kim JS, Hwang B-H, Lee C-S. A Co-culture of Staphylococcus epidermidis and Staphylococcus aureus in a Monodisperse Droplet to Investigate Microbial Interaction at Defined Microenvironment. Korean Journal of Chemical Engineering. 2025;42(10):2355–71.

34. Tan JY, Wang S, Dick GJ, Young VB, Sherman DH, Burns MA, et al. Co-cultivation of microbial sub-communities in microfluidic droplets facilitates high-resolution genomic dissection of microbial ‘dark matter’. Integrative Biology. 2020;12(11):263–74.

35. Huang C, Jiang Y, Li Y, Zhang H. Droplet detection and sorting system in microfluidics: a review. Micromachines. 2022;14(1):103.

36. Jiang J, Yang G, Ma F. Fluorescence coupling strategies in fluorescence-activated droplet sorting (FADS) for ultrahigh-throughput screening of enzymes, metabolites, and antibodies. Biotechnology advances. 2023;66:108173.

37. Panwar J, Autour A, Merten CA. Design and construction of a microfluidics workstation for high-throughput multi-wavelength fluorescence and transmittance activated droplet analysis and sorting. Nature protocols. 2023;18(4):1090–136.

38. Li M, Liu H, Zhuang S, Goda K. Droplet flow cytometry for single-cell analysis. RSC advances. 2021;11(34):20944–60.

39. Dsouza A, Taylor D, Parmenter C, Hand RA, Brettschneider J, Unnikrishnan M, et al. Dynamic bacterial growth modulation in structurally distinct and functionally tuneable agarose hydrogels. Communications Materials. 2026;7(1):57. doi: 10.1038/s43246-025-01067-9.

40. Duarte JM, Barbier I, Schaerli Y. Bacterial Microcolonies in Gel Beads for High-Throughput Screening of Libraries in Synthetic Biology. ACS Synthetic Biology. 2017;6(11):1988–95. doi: 10.1021/acssynbio.7b00111.

41. Kandemir N, Vollmer W, Jakubovics NS, Chen J. Mechanical interactions between bacteria and hydrogels. Scientific reports. 2018;8(1):10893.

42. Schultz KM, Anseth KS. Monitoring degradation of matrix metalloproteinases-cleavable PEG hydrogels via multiple particle tracking microrheology. Soft matter. 2013;9(5):1570–9.

43. Lutolf MP, Lauer-Fields JL, Schmoekel HG, Metters AT, Weber FE, Fields GB, et al. Synthetic matrix metalloproteinase-sensitive hydrogels for the conduction of tissue regeneration: engineering cell-invasion characteristics. Proceedings of the National Academy of Sciences. 2003;100(9):5413–8.

44. Kim Y-M, Kiso Y, Muraki T, Kang M-S, Nakai H, Saburi W, et al. Novel dextranase catalyzing cycloisomaltooligosaccharide formation and identification of catalytic amino acids and their functions using chemical rescue approach. Journal of Biological Chemistry. 2012;287(24):19927–35.

45. Staat RH, Schachtele CF. Evaluation of dextranase production by the cariogenic bacterium Streptococcus mutans. Infection and Immunity. 1974;9(2):467–9.

46. Bassis CM, Tang AL, Young VB, Pynnonen MA. The nasal cavity microbiota of healthy adults. Microbiome. 2014;2(1):27.

47. Darwish MS, Khalil WA, Hassan MA, Moussa M, Abou-Zeid NA, Abdelnour SA, et al. Exploring the Role of E. Coli Nissle 1917 Postbiotics as Antimicrobial and Antioxidant Agents for Enhancing Buffalo Sperm Quality. Probiotics and Antimicrobial Proteins. 2026:1-24.

48. Fang K, Jin X, Hong SH. Probiotic Escherichia coli inhibits biofilm formation of pathogenic E. coli via extracellular activity of DegP. Scientific reports. 2018;8(1):4939.

49. Baret J-C, Miller OJ, Taly V, Ryckelynck M, El-Harrak A, Frenz L, et al. Fluorescence-activated droplet sorting (FADS): efficient microfluidic cell sorting based on enzymatic activity. Lab on a Chip. 2009;9(13):1850–8.

50. Sukovich DJ, Kim SC, Ahmed N, Abate AR. Bulk double emulsification for flow cytometric analysis of microfluidic droplets. Analyst. 2017;142(24):4618–22.

51. Eun Y-J, Utada AS, Copeland MF, Takeuchi S, Weibel DB. Encapsulating bacteria in agarose microparticles using microfluidics for high-throughput cell analysis and isolation. ACS chemical biology. 2011;6(3):260–6.

52. Li P, Müller M, Chang MW, Frettlöh M, Schönherr H. Encapsulation of autoinducer sensing reporter bacteria in reinforced alginate-based microbeads. ACS applied materials & interfaces. 2017;9(27):22321–31.

53. Ochoa A, Gastélum G, Rocha J, Olguin LF. High-throughput bacterial co-encapsulation in microfluidic gel beads for discovery of antibiotic-producing strains. Analyst. 2023;148(22):5762–74.

54. Cruz-Acuña R, Quirós M, Farkas AE, Dedhia PH, Huang S, Siuda D, et al. Synthetic hydrogels for human intestinal organoid generation and colonic wound repair. Nature cell biology. 2017;19(11):1326–35.

55. Lin L, Zhu J, Kottke-Marchant K, Marchant RE. Biomimetic-engineered poly (ethylene glycol) hydrogel for smooth muscle cell migration. Tissue Engineering Part A. 2014;20(3-4):864–73.

56. Habibi N, Mustafa AS, Khan MW. Composition of nasal bacterial community and its seasonal variation in health care workers stationed in a clinical research laboratory. PLoS One. 2021;16(11):e0260314. doi: 10.1371/journal.pone.0260314.

57. Aggarwal D, Bellis KL, Blane B, de Goffau MC, Wagner J, Ng DYK, et al. Large-scale characterisation of the nasal microbiome redefines Staphylococcus aureus colonisation status. Nature Communications. 2025;16(1):10415. doi: 10.1038/s41467-025-66564-4.

58. Jha RK, Strauss CE. Smart microbial cells couple catalysis and sensing to provide high-throughput selection of an organophosphate hydrolase. ACS Synthetic Biology. 2020;9(6):1234–9.

59. Alshareedah I, Kumar A. Multiphasic droplet microfluidics platform for controlled bacteria and mammalian cell co-culture. Lab on a Chip. 2026;26(8):2473–85.

